# A brassinosteroid receptor kinase is required for sex determination in the homosporous fern *Ceratopteris richardii*

**DOI:** 10.1101/2024.10.09.617452

**Authors:** Katelin M. Burow, Xi Yang, Yun Zhou, Brian P. Dilkes, Jennifer H. Wisecaver

**Affiliations:** 1Department of Biochemistry, Purdue University, _West Lafayette_, Indiana, USA; 2Purdue Center for Plant Biology, Purdue University, _West Lafayette_, Indiana, USA; 3Department of Botany and Plant Pathology, Purdue University, _West Lafayette_, Indiana, USA

## Abstract

Most ferns, unlike all seed plants, are homosporous and produce sexually undifferentiated spores. Sex in many homosporous ferns is environmentally regulated by the biosynthesis of antheridiogens. Secreted antheridiogen from female or hermaphrodite gametophytes is perceived by undetermined gametophytes which respond by developing as male. In the fern *Ceratopteris richardii* (Ceratopteris), several *hermaphroditic* (*her*) mutants develop as hermaphrodites even in the presence of antheridiogen. A set of five mutant alleles are tightly linked to *her7*, a locus of unknown molecular identity. The immense size of the Ceratopteris genome has made efficient identification of the underlying genes a challenge. Here, we used bulked segregant RNA-seq (BSR-Seq) to map two *her* mutants (*her7-14* and *her7-19*) to a 16 Mbp interval on chromosome 29 of the Ceratopteris genome. A brassinosteroid receptor-like kinase (BRL) within this interval encoded a deletion mutation in *her7-14* and a missense mutation in *her7-19*. Three other linked *her* mutants (*her7-1*, *her7-11*, and *her7-15*) encoded missense mutations in the same gene, which we name HER7, providing strong support for HER7 being required for antheridiogen-mediated sex determination in Ceratopteris. HER7-GFP fusion protein is localized in the plasma membrane and cytoplasm, consistent with its predicted function as a brassinosteroid receptor kinase. RNA-seq analysis of gene expression shows that numerous transcripts encoding enzymes, including those for brassinosteroid biosynthesis, accumulate in hermaphrodites compared to male gametophytes. The antheridiogen of Ceratopteris is currently unknown and our finding that a BRL mutant is insensitive to antheridiogen suggests that antheridiogen in Ceratopteris may be a brassinosteroid. Additional work is required to fully resolve the sex determination pathway and determine the chemical composition of antheridiogen in this species. Our work demonstrates we can identify the molecular basis of mutants in large genomes, such as those in homosporous ferns, which can be further applied to the extensive remaining collection of mutants in Ceratopteris.

## INTRODUCTION

Ferns are the second most diverse group of vascular plants after angiosperms (PPG 2016). Most ferns are homosporous, producing a single spore type at meiosis (Moran 2004). This contrasts with heterosporous plants (such as seed plants and a minority of fern species) that produce mega- and microspores that develop into predetermined female and male gametophytes, respectively. In homosporous species, the sex of the haploid gametophyte (which can be male, female, or hermaphroditic depending on the system) is instead determined by a programmed response to environmental factors following spore germination. A common method of environmental regulation of sex in homosporous ferns is the production of pheromones known as antheridiogens that influence the sex determination of neighboring gametophytes (Hornych et al. 2021).

Antheridiogens are signaling molecules released by female and hermaphroditic fern gametophytes that promote the development of male traits including antheridia (sperm- forming structures) while simultaneously suppressing the development of female traits including the meristem and archegonia (egg-forming structures) (Schneller 2008). For example, in the model fern *Ceratopteris richardii* (hereafter Ceratopteris), a spore grown in isolation will always develop as a hermaphrodite that produces and secretes <u>a</u>ntheridiogen of <u>Ce</u>ratopteris (A_CE_) into its surroundings. In contrast, nearly all spores (>95%) that germinate in the presence of A_CE_ will develop as males (Banks et al. 1993). The two sex types are morphologically distinct; the mature hermaphrodite prothallus has archegonia, antheridia, and a multicellular meristem (Figure 1A), whereas most cells of the male prothallus differentiate as antheridia and do not form a meristem (Figure 1B).

**Figure 1:**
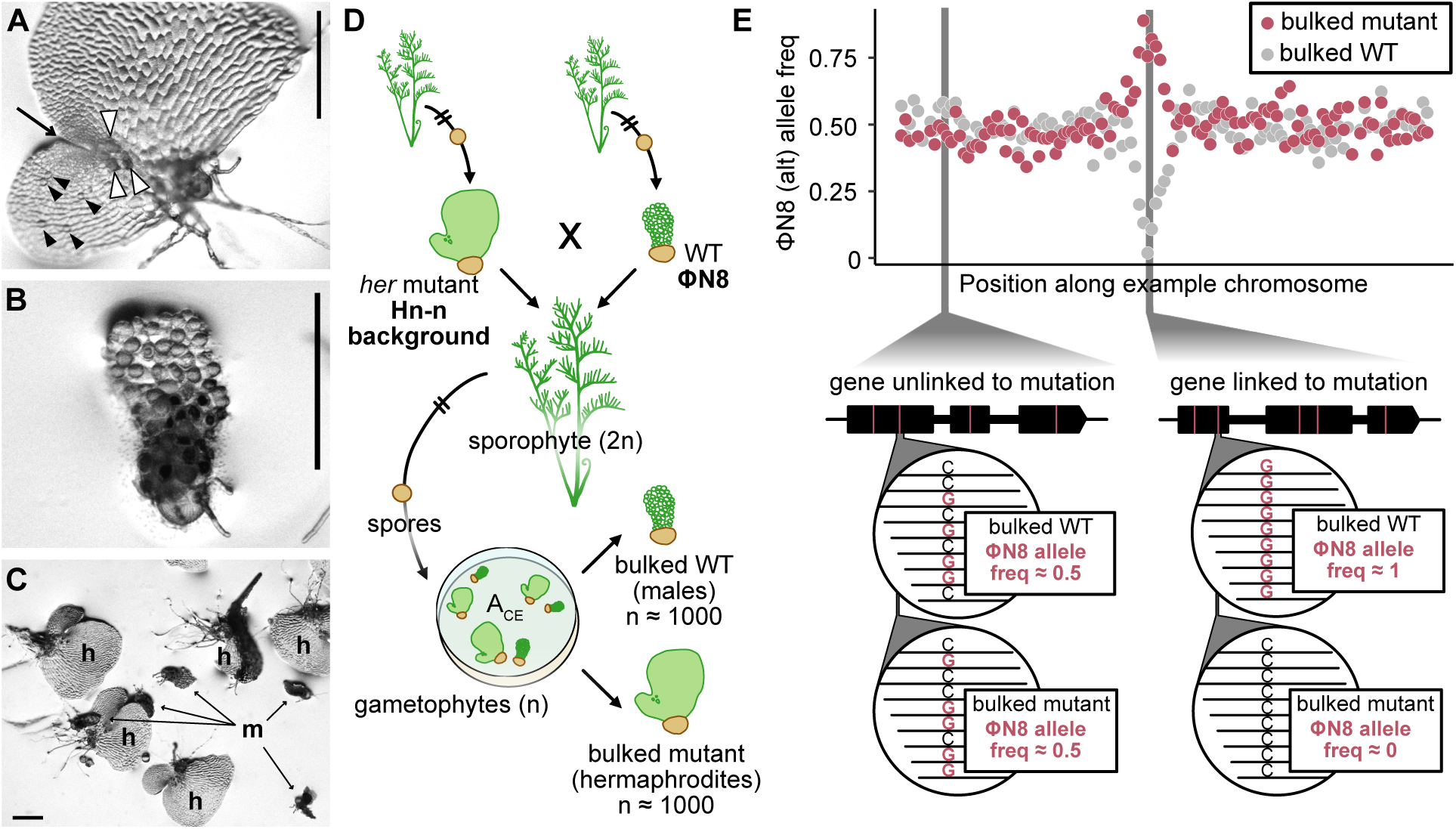
Overview of Ceratopteris gametophyte morphology and BSR-Seq pipeline. (A) Hermaphrodite gametophyte showing location of archegonia (white arrows), antheridia (black arrows), and notched meristem (thin arrow). (B) Male gametophyte consisting almost entirely of antheridia. (C) Hn-n gametophytes on FM plate showing clear phenotypic difference between hermaphrodites (h) and males (m). Scale bars: 500 μm. (D) Workflow for creating hybrid crosses. (E) Overview of methodology for identifying genome region linked to causative SNPs.

A_CE_ is secreted by the hermaphrodite after it becomes insensitive to its effects (Banks et al. 1993). Males require constant exposure to A_CE_ to remain male. Once removed from A_CE_-containing media, cells within the male divide and differentiate into a hermaphroditic prothallus (Banks et al. 1993).

Multiple antheridiogen systems exist in ferns. The A type antheridiogens were first identified in *Pteridium aquilinum* by Döpp (1950) and similar antheridiogens are produced by many species within the Polypodiales (Schneller 2008; Hornych et al. 2021). B type antheridiogens are produced by species within the Schizaeales, including *Anemia* and *Lygodium.* Lastly, the C type antheridiogen (e.g., A_CE_) is thus far limited to members of the *Ceratopteris* genus. While the structures of A and C type antheridiogens are unknown, all characterized B type antheridiogens are gibberellins (GAs) (Corey et al. 1986; Takeno et al. 1989; Yamauchi et al. 1996; Yamane 1998; Kurumatani et al. 2001). Given the distribution of antheridiogen systems across the fern phylogeny, it is unclear when and how these different antheridiogen types first evolved and diversified (Hornych et al. 2021). Yet, some evidence suggests GA-based signaling may also be involved in Ceratopteris sex determination, as it is in B type antheridiogen-containing species. The presence of GA biosynthetic inhibitors AMO-1618 (2’-isopropyl-4’- (trimethylammoniumchloride)-5’-methylphenylpiperidine-1-carboxylate) and CCC (2- chloroethyl trimethylammonium chloride) in growth media slightly reduces the proportion of males in a gametophyte population (Warne and Hickok 1989). Moreover, abscisic acid (ABA) has many characterized antagonistic interactions with GA in angiosperms (Li et al. 2002; Achard et al. 2006; Finch-Savage and Leubner-Metzger 2006) and completely blocks the A_CE_ response in Ceratopteris (Hickok 1983).

Despite the unknown chemical identity of A_CE_, Ceratopteris has been extensively used to study the genetics of antheridiogen-based sex determination for decades. Ceratopteris traits conducive to genetic research include the clear sexual dimorphism of its two sexes (Figure 1C), easily crossable nature, and short life cycle (Hickok et al. 1987, 1995; Eberle et al. 1995; Chatterjee and Roux 2000). Because gametophytes are haploid, dominant and recessive mutants are easy to select from a population of mutagenized single-celled spores. The abundance of spores produced by a single homozygous plant allows suppressors of characterized mutations to be readily identified in only one generation (e.g., Banks 1994). As a result, dozens of sex determination mutants with diverse developmental phenotypes have been genetically characterized in Ceratopteris (Warne et al. 1988; Banks 1994, 1997; Eberle and Banks 1996; Strain et al. 2001). Among them are the *hermaphroditic* (*her*) mutants that develop as hermaphrodites even in the presence of A_CE_. The first *her* mutant was isolated over 30 years ago (Warne et al. 1988), yet the molecular identities of these mutant genes remain unknown. The large size of the Ceratopteris genome (1n = 9.6 Gb) has hindered traditional genetic methods for gene identification. Here, we leveraged the recent sequencing of the Ceratopteris genome (Marchant et al. 2022) combined with the throughput and cost effectiveness of bulk segregant analysis by RNA-seq (BSR-Seq) to clone the first mutant gene required for sex determination in Ceratopteris.

## RESULTS

### *Mutant mapping by BSR-seq identifies HER7*, a receptor-like kinase required for antheridiogen perception

All *her* sex determination mutants used in our analysis were previously generated in the same Hn-n genetic background as the Ceratopteris reference genome (Banks 1994; Marchant et al. 2022). To enable mapping of the *her* mutations, we first sequenced the transcriptome of a divergent Ceratopteris genotype, ΦN8, from Nicaragua (Hickok et al. 1995). Single nucleotide polymorphisms were identified by mapping Illumina RNA-seq reads from ΦN8 to the Hn-n Ceratopteris reference genome, which yielded a total of 40,328 high-quality, single nucleotide variants distributed across the genome. This provided sufficient genotypic diversity within coding regions to perform informative hybrid crosses.

Previously, *her7*, *her10*, *her11*, *her14*, *her15*, and *her19* were demonstrated to be linked, and possibly represent allelic mutants (Eberle and Banks, 1996). These mutants produced no males on A_CE_, with the exception of *her14* which encodes a weak allele and a small number of individuals developed into males. Spore progeny from F1 *her14/+* Hn-n × ΦN8 and *her19/+* Hn-n × ΦN8 hybrids were grown in the presence of A_CE_. Developing gametophytes segregated for the *her* mutations, resulting in hermaphrodites containing the *her* mutation and males containing the wild type ΦN8 genotype at the causative locus (Figure 1D). Bulked hermaphrodite and male samples of ∼1000 individuals were sequenced by BSR-seq, and the frequency of the ΦN8 allele (*i.e.,* read depth of the ΦN8 allele divided by total read depth) was determined. In hermaphrodite and male bulked samples, the distribution of the ΦN8 genotype fluctuated around the 0.5 expected frequency across most of the genome. At the region surrounding the *her* mutations, the frequency of the ΦN8 allele approached zero in the hermaphrodite samples and 100% in the male samples (Figure 1E). Both the *her14* and *her19* mutations mapped to the same region on chromosome 29, as expected, overlapping in a 16 Mbp window (Figure 2A). This window contained 77 genes (Figure 2B). Due to the low sequencing depth of our *her* x ΦN8 BSR-Seq samples, we analyzed a *her19* RNA-seq dataset from an uncrossed mutant (Geng et al. 2021) to identify candidate causal mutations for the *her19* phenotype.

**Figure 2:**
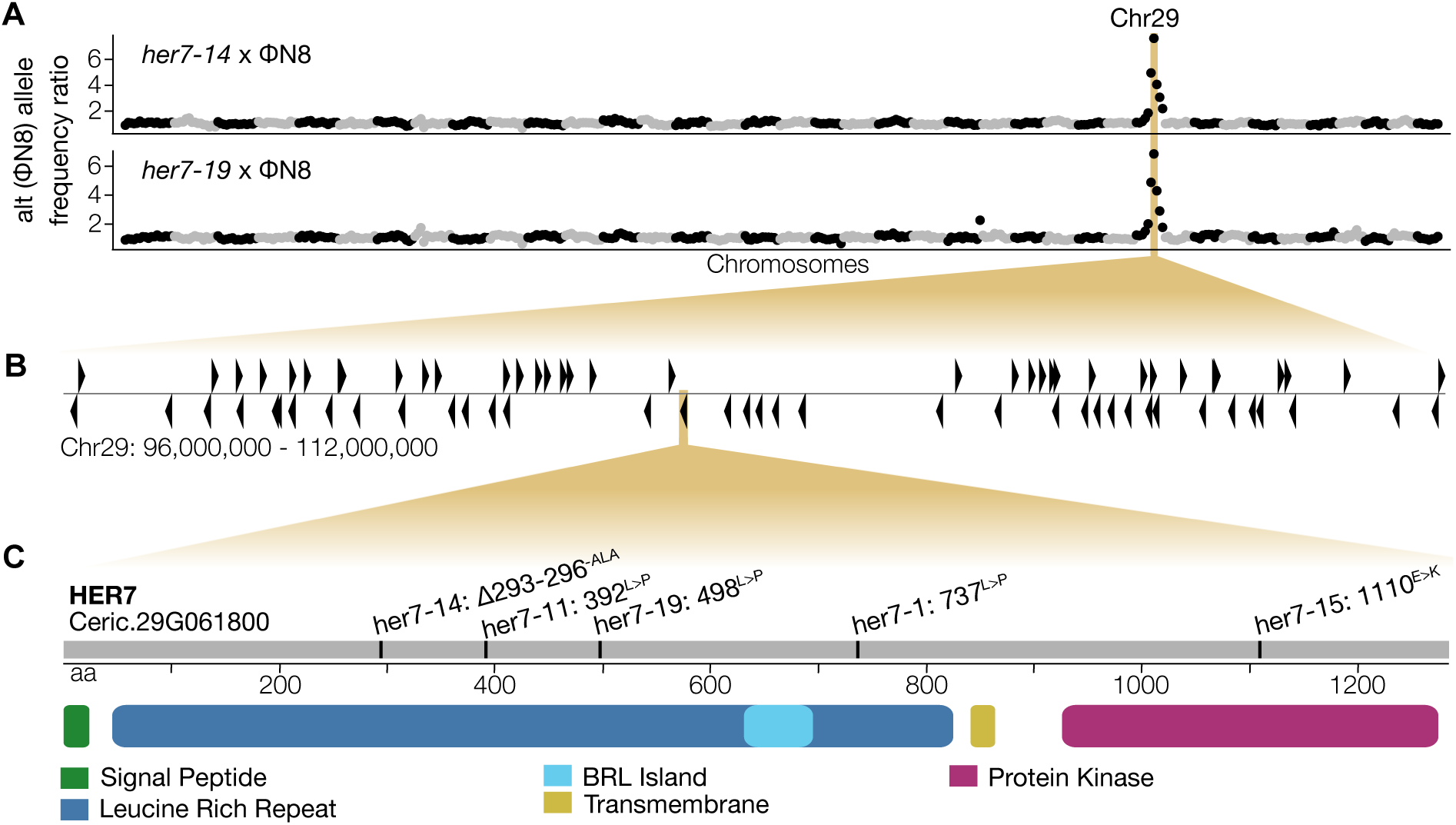
BSR-Seq mapping results. (A) Distribution of the alternative (N8) allele across the Ceratopteris genome in two *her* × ΦN8 crosses. (B) Distribution of protein-coding genes in mapped window. (C) Nonsynonymous mutations were identified in *HER7* in five independent *her* mutants. Protein domain boundaries were identified using the InterPro online search tool.

Within the mapped region, candidate missense variants were detected in two genes: Ceric.29G061800, encoding a putative receptor-like kinase; and Ceric.29G060400, a gene of unknown function containing multiple armadillo-like α-helix repeats (Table S1). All five variants in Ceric.29G060400 had allele frequencies less than 100% and were likely the result of poor read mapping in the repetitive regions of the gene. Therefore, we chose to focus our investigation on the putative mutation in Ceric.29G061800, which had an allele frequency of 100% in the *her19* mutant and resulted in the conversion of a leucine to a proline (L498P) within the extracellular leucine-rich repeat (LRR) domain of the protein (Figure 2C). A search for conserved domains in addition to the LRR domain identified a signal peptide, transmembrane domain, and C-terminal kinase domain (Figure 2C). PCR sequencing of Ceric.29G061800 in the mutants *her7*, *her11*, *her14*, and *her15* identified novel nonsynonymous mutations in this gene in each of the mutants (Figure 2C). For clarity, we name this locus *her7* after the first mutant identified in this linkage group and designate the alleles based on the mutant number given at the time of isolation (i.e., *her7-1*, *her7-11*, *her7-14*, *her7-15,* and *her7-19*). The *her7-1* and *her7-11* mutations were leucine to proline conversions, L737P and L392P respectively, within the LRR domain. The weak *her7-14* allele resulted from a three-amino acid deletion (-ALA Δ293-296) in the same LRR domain. The fourth mutation, *her7-15,* caused a glutamic acid to lysine conversion (E1110K) within the intracellular protein kinase domain (Figure 2C) resulting in the same change at the homologous position as the *bri1-703* allele from Arabidopsis (Sun et al. 2017). Five independent nonsynonymous mutations at the same locus within the mapping window demonstrates that Ceric.29G061800 encodes *HER7* and is required for antheridiogen perception or downstream sex determination in Ceratopteris.

#### HER7 is closely related to BRI1 brassinosteroid receptors of seed plants

A phylogenetic analysis of the HER7 protein sequence places it within the BRASSINOSTEROID INSENSITIVE 1 (BRI1)-like family of leucine-rich repeat receptor-like kinases (Figure 3A). In the model angiosperm *Arabidopsis thaliana* (hereafter Arabidopsis), BRI1 is the primary receptor required for brassinosteroid (BR) signaling (Noguchi et al. 1999); binding of BRs to the extracellular domain of BRI1 activates the intracellular kinase domain and relays the signal to the cytoplasmic side of plasma membrane (He et al. 2000). In our phylogeny of BRI1-like receptors (BRLs), HER7 groups with strong support inside a homosporous fern-specific clade that is sister to the BRI1-containing clade of seed plants (Figure 3A). An additional paralog in Ceratopteris (HER7-like, Ceric.21G054300) is also present in the HER7 clade (Figure 3A). Notably, this clade has been lost in all heterosporous ferns sequenced to date (Figure 3A).

**Figure 3:**
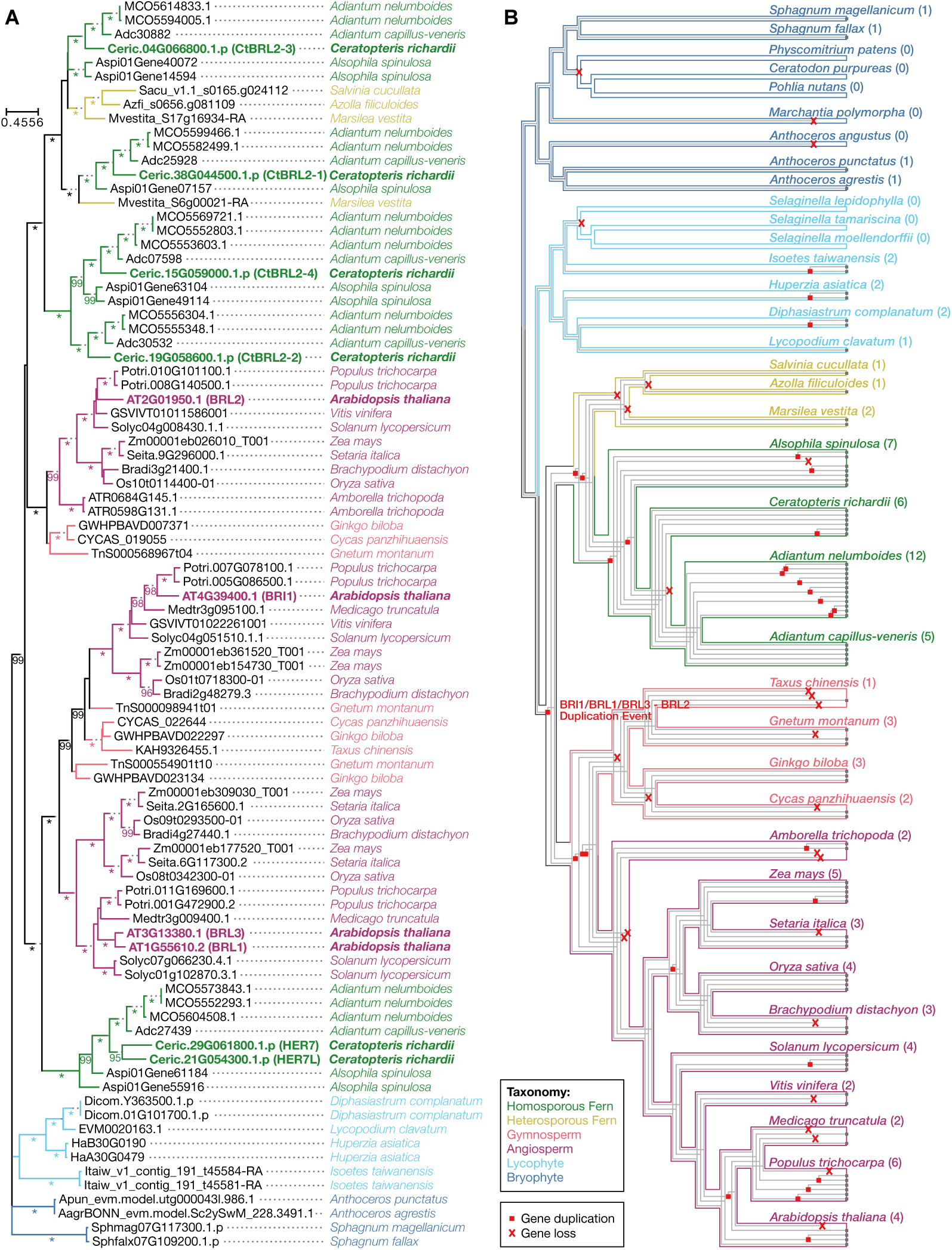
Phylogenetic analysis of the BRL gene family in plants. (A) GeneRax- optimized maximum likelihood phylogeny. Arabidopsis and Ceratopteris homologs are bolded. Nodes with ultrafast bootstrap support > 95 are labeled along the preceding branch; ultrafast bootstrap support of 100 is indicated by an asterisk (*); nodes that lack branch support are unlabeled. (B) GeneRax-inferred patterns of gene duplications and loss of BRL homologs on the species tree. Total BRL gene copies present in each species is indicated by the number in parenthesis.

Four additional BRLs were previously identified in Ceratopteris (CtBRL2-1, CtBRL2-2, CtBRL2-3, and CtBRL2-4; Zheng et al. 2022), all four of which group with strong support inside a larger fern clade comprised of sequences from both homosporous and heterosporous species (Figure 3A). This second fern clade groups sister to the BRL2- containing clade of seed plants; however, this association lacks branch support (Figure 3A). We performed a topology test to evaluate the support for the presence of two distinct fern clades in the BRL gene family and found that the ML phylogeny presented in Figure 3A was significantly better than an alternative topology that forced all fern sequences to be monophyletic (approximately unbiased test = 0.005). Taken together, our phylogenetic analysis provides strong support that the sequences in the HER7 clade are the most closely related fern homologs to the BRI1-containing clade in seed plants which encodes all BR receptors characterized to date.

We performed a gene tree-species tree reconciliation using GeneRax (Morel et al. 2020) to model patterns of gene duplication and loss across the BR1/BRL phylogeny.

Our analysis indicates that BR1/BRL gene family expansion first occurred via a duplication event in the last common ancestor (LCA) of euphyllophytes (ferns and seed plants) giving rise to the BRI1 and BRL2 clades (Figure 3B). Subsequent lineage-specific duplication events further diversified the gene family (Figure 3B). It is notable that *HER7* is absent in all three heterosporous ferns in our analysis. The reconciliation analysis suggests that *HER7* was lost in these heterosporous species (Figure 3B) and is correlated with the transition to heterospory in these taxa.

#### HER7 is localized to the plasma membrane

To examine the subcellular localization of HER7, we generated an expression cassette directing the expression of a GFP-HER7 fusion protein under the control of the Cauliflower Mosaic Virus 35S promoter (*35S::GFP-HER7)*. We compared the fluorescence localization in transiently transformed cells to a GFP-only control (*35S::GFP*). Confocal imaging revealed that in cells transiently expressing GFP alone (Figure 4A,B), GFP signals were ubiquitously detected throughout the cell, including the nucleus, cytosol, and plasma membrane. In contrast, GFP-HER7 signals were localized specifically to the plasma membrane and cytoplasm (Figure 4C,D), consistent with the predicted function of HER7 as a receptor kinase.

**Figure 4:**
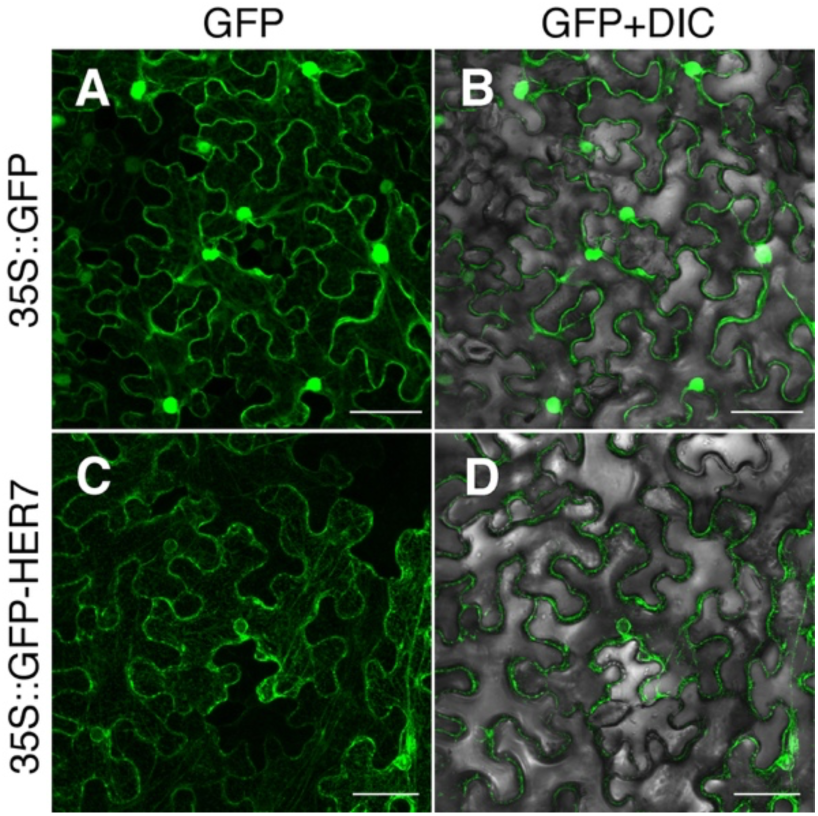
Subcellular localization of HER7. Confocal microscopy images of *N. benthamiana* cells transiently expressing either GFP only (A, B) or the GFP-HER7 fusion protein (C, D). GFP channel (A, C) and merged images of GFP and DIC channels (B, D). Scale bar: 50 μm. At least three samples were imaged under the same conditions, showing comparable results.

### Genes for brassinosteroid and gibberellin signaling and biosynthesis are differentially expressed between male and hermaphroditic gametophytes

We identified differentially expressed genes (DEG) by comparing transcript accumulation by RNAseq of *her7-19,* wild type hermaphrodite, and wild type male gametophytes. A significant number of genes were differentially expressed in all three pairwise comparisons using a false discovery rate of 0.1 and absolute log_2_ fold change (LFC) cutoff of 1.0. The greatest number of DEGs were observed in comparisons between male and hermaphrodite gametophytes; n = 6,551 in the male vs wild type hermaphrodite comparison (Figure 4A) and n = 8,481 in the male vs mutant *her7-19* comparison (Figure 4B). Fewer genes (n = 1,466) were differentially expressed in the *her7-19* versus wild type hermaphrodite comparison with the majority of DEGs in this category displaying reduced expression in *her7-19* (Figure 4C).

To determine the functional categories that were significantly enriched in DEGs, we used the Kyoto Encyclopedia of Genes and Genomes (KEGG) (Kanehisa et al. 2014) to assign Ceratopteris genes to overlapping, higher-order biological pathways (Table S2). Anabolic metabolism and essential pathways were broadly upregulated in hermaphrodites as compared to males (BH adjusted p-value < 0.05; Figure 4D).

Specialized metabolism biosynthesis pathways including those for the production of phenylpropanoids and terpenoids were also enriched in these same DEG sets. In contrast, KEGG pathways for MAPK and signal transduction as well as cellular processes including cell growth, cell death were all significantly enriched in DEGs with higher expression in the male samples compared to hermaphrodites (BH adjusted p- value < 0.05; Figure 4D). Cell motility was also enriched in DEGs with higher expression in the males, as expected due to the high proportion of cells that were motile sperm.

These signal transduction and cellular processes KEGG pathways were also enriched in DEGs with lower expression in *her7-19* mutant hermaphrodites compared to wild type hermaphrodites (Figure 4D). No KEGG categories were significantly enriched in the few DEGs with higher expression in *her7-19* mutant hermaphrodites compared to wild type (Figure 4D).

We next compared the expression of several gene families known to be involved in BR signaling and biosynthesis (Table S3, Table S4). *HER7* itself was not significantly differentially expressed in any comparison; however, three *BRI*/*BRL* paralogs including *HER7-like*, *CtBRL2-1,* and *CtBRL2-3* transcripts were less expressed in males compared to both wild type and *her7-19* hermaphrodites (BH adjusted p-value ≤ 0.0189; LFC ≤ -2.00; Figure 5). Ceratopteris encodes two homologs of *BRI1-ASSOCIATED RECEPTOR KINASE1* (*BAK1*), and four homologs of *BRASSINOSTEROID- SIGNALING KINASE1* (*BSK1*) both of which are required for BR signaling in Arabidopsis (Li et al. 2002; Shi et al. 2013). One copy from each of these gene families was less expressed in males compared to hermaphrodites (BH adjusted p-value ≤ 4.74 x 10^-3^; LFC ≤ -1.49; Figure 5). Ceratopteris also encodes five homologs of *BRI1 SUPPRESSOR1* (*BSU1*) a phosphatase involved in BR signaling in Arabidopsis (Ryu et al. 2010), none of which were significantly DE in our analysis (Table S4; Figure 5).

**Figure 5:**
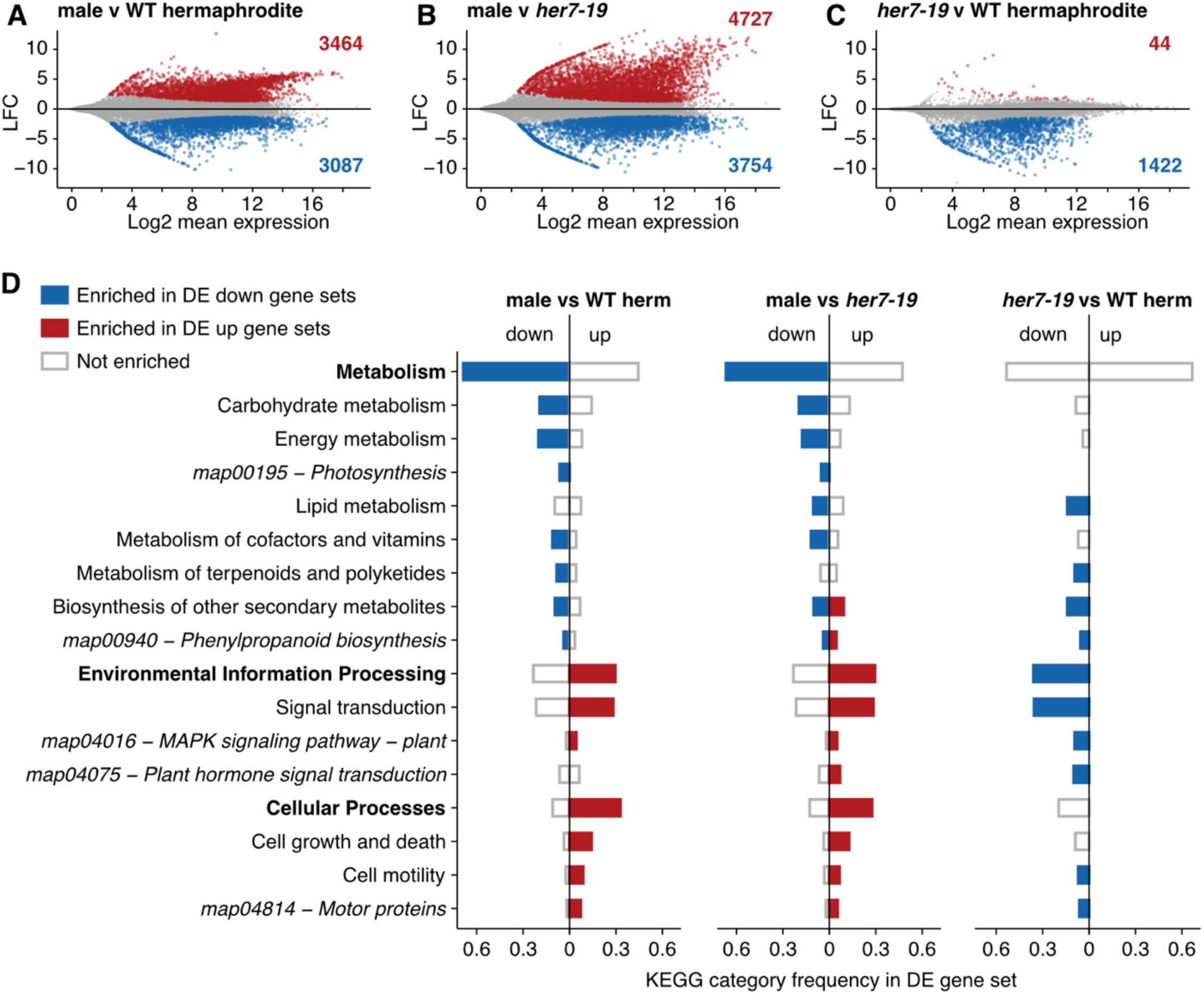
Differential gene expression in male and hermaphrodite gametophytes. (A-C) MA-plots showing differentially expressed genes with increased abundance (red points and numbers) or decreased abundance (blue points and numbers) in the first condition versus the second condition using an adjusted p-value < 0.1 and absolute LFC > 1.0. (D) Significantly enriched KEGG pathways (italics), pathway categories (unbolded), higher order category (bolded). Bar height indicates frequency of the annotation in DE gene sets.

Cytochrome P450s encoded by the CYP90 gene family are required for BR biosynthesis in angiosperms (Chung and Choe 2013; Vriet et al. 2013). We identified two homologs of CYP90B, both of which were less expressed in males compared to hermaphrodite gametophytes (BH adjusted p-value ≤ 5.53 x 10^-3^; LFC ≤ -3.25; Figure 5). We identified a single homolog of CYP90A, which was not significantly DE in any comparison (Table S4; Figure 5). The one CYP90C homolog in Ceratopteris was decreased in its transcript accumulation in males compared to hermaphrodites (BH adjusted p-value ≤ 6.10 x 10^-7^; LFC ≤ -3.04; Figure 5). CYP85A1 acts late in the pathway and is responsible the production of castasterone and brassinolide, the two dominant bio-active BRs in angiosperms (Kim et al. 2008). No CYP85A1 homologs were detected in any ferns in our analysis (Table S3), including Ceratopteris, consistent with the lack of brassinolide and very low levels of castasterone detected thus far in ferns (Yokota et al. 2017). We also identified four homologs of *DEETIOLATED2* (*DET2*), a steroid 5-alpha-reductase involved in sterol biosynthesis and required for BR biosynthesis in angiosperms (Fujioka et al. 1997). Contrasting the pattern observed in the CYP90 homologs, one *DET2* gene in Ceratopteris had significantly higher expression in males compared to hermaphrodites (BH adjusted p-value ≤ 5.65 x 10^-22^; LFC ≥ 3.17; Figure 5). This indicates that there is a strong upregulation of the CYP90 BR biosynthetic enzymes in hermaphrodites consistent with, but not demonstrating that, a brassinosteroid could act as A_CE_.

GAs were previously implicated in sex determination in Ceratopteris through the use of biosynthetic inhibitors (Hickok 1983; Warne and Hickok 1989). We investigated the expression pattern of genes involved in GA signaling and biosynthesis in males, hermaphrodites and *her7-19* mutants (Table S3, Table S4). We identified four homologs of the receptor GIBERELLIN INSENSITIVE1 (GID1), which binds GA and is involved in GA hormone signaling (Ueguchi-Tanaka et al. 2005). Transcripts from two GID1 homologs were accumulated in males compared to hermaphrodites (BH adjusted p-value ≤ 4.55 x 10^-3^; LFC ≥ 2.45; Figure 5). Two homologs of the DELLA-domain encoding transcription factors that repress GA signaling (Hussain and Peng 2003) are present in the Ceratopteris genome, neither of which were significantly differentially expressed in our analysis (Table S4; Figure 5). We also identified the Ceratopteris homologs of seven gene families involved in GA biosynthesis in Arabidopsis: *ENT-COPALYL DIPHOSPHATE SYNTHETASE* (*CPS*), *ENT-KAURENE SYNTHASE* (*KS*), *ENT-KAURENE OXIDASE* (*KO*), *ENT-KAURENOIC ACID HYDROXYLASE* (*KAO*), *GIBBERELLIN 2-OXIDASE* (*GA2OX*), *GIBBERELLIN 3-OXIDASE* (*GA3OX*), and *GIBBERELLIN 20-OXIDASE* (*GA20OX*) (Sun and Kamiya 1994; Helliwell et al. 1998, 2001; Yamaguchi et al. 1998; Thomas et al. 1999; Hedden and Phillips 2000). No *GA2OX* homologs were identified in ferns, including Ceratopteris (Table S3). The *CPS* and *KS* gene families formed one large orthogroup (Table S3), and three of the five *CPS*-*KS* homologs in Ceratopteris were DE with two transcripts decreased and one increased in accumulation in males as compared to hermaphrodites (Table S4; Figure 5). We identified a single *KAO* homolog which was less expressed in males compared to hermaphrodites (BH adjusted p-value ≤ 2.58 x 10^-2^; LFC ≤ -2.94; Figure 5).

Ceratopteris also possesses one homolog each of *KO* and *GA3OX*, neither of which were differentially expressed in our analysis (Table S4; Figure 5). The six Ceratopteris *GA20OX* homologs displayed the most complicated pattern of expression and included one copy that was differentially expressed between the mutant *her7-19* and wild type hermaphrodites (Table S4; Figure 5). Taken together, *HER7* being a brassinosteroid receptor-like gene, the upregulation of BR biosynthetic enzymes in hermaphrodites, and the lack of a clear pattern in the expression of GA biosynthetic machinery, raises the possibility that A_CE_ is a BR.

## DISCUSSION

The identification of *HER7*, a gene encoding a BR receptor-like kinase, as the causal locus for the hermaphroditic mutant phenotype in five *her7*-linked mutants raises several exciting new questions about plant sex determination and antheridiogen signaling. HER7 is orthologous to BRI1, the primary BR receptor in angiosperms, and the two proteins share the same domain structure. In BRI1, the leucine-rich repeat (LRR) domain binds BRs (She et al. 2011). Three *her* mutants (*her7-1, her7-11*, and *her7-19*) encode leucine to proline missense mutations in the LRR domain. A 9- nucleotide deletion mutant (*her7-14)* was also mapped to the LRR domain. The *her7-14* mutant encodes a weak allele resulting in a sex ratio with ∼3% males (Banks 1994). The *her7-15* mutation changes a negatively charged glutamic acid to a positively charged lysine in the activation domain of the C-terminal kinase. This is the identical substitution at the homologous position that causes the *bri1-703* allele, which has been biochemically characterized in Arabidopsis and shown to disrupt the function of the protein kinase of BRI1 (Sun et al. 2017). In *her7* mutants, gametophytes develop as hermaphrodites even in the presence of A_CE_. This demonstrates that *HER7* is required for A_CE_ to induce or maintain male development and strongly suggests that perception of a BR is required for male development in Ceratopteris.

Brassinosteroids have many diverse regulatory functions in angiosperms including influencing cell elongation and division, vascular differentiation, photomorphogenesis, organ identity, meristem identity, seed germination, immunity, and stomata development (Cheon et al. 2010; Hartwig et al. 2011; Kim et al. 2012; Makarevitch et al. 2012; Wang et al. 2012; Best et al. 2016; Yang et al. 2018; Furuya et al. 2024). Less is known about the function of brassinosteroids in lineages outside of angiosperms. However, plants as distant from angiosperms as the lycophyte Selaginella encode *CYP90* genes and will elongate in response to BR application (Cheon et al. 2013). BR metabolites have been identified in ferns, including the bioactive castasterone, but the ability of ferns to produce brassinolide, the most bioactive BR compound in Arabidopsis, is unclear (Yokota et al. 2017; Fernández et al. 2021). The endogenous levels of known bioactive brassinosteroids reported in ferns were far lower than that typically reported in angiosperms (Yokota et al. 2017; Fernández et al. 2021). These methods only detect known brassinosteroids, and as a result, differences in the pathway intermediates or accumulated active forms would result in a failure to detect or measure the bioactive BRs in Ceratopteris. Additional work to characterize the brassinosteroids made by non-flowering plants, including Ceratopteris, is necessary to clarify this.

Disrupting the *HER7* locus does not result in a mutant sporophyte phenotype (Banks 1994). This indicates that HER7 is not required for BR signaling in the sporophyte.

However, exogenously applied brassinolide increased the rate of Ceratopteris sporophyte growth (Zheng et al. 2022). Our phylogenetic analysis identified two copies of BRI1 orthologs in Ceratopteris, HER7 and HER7-like (Figure 3). It is possible that following gene duplication, the function of these two genes diverged with HER7-like being responsible for BR perception in the sporophyte and HER7 specialized for sex determination. However, BRI1 orthologs were conspicuously absent in all heterosporous ferns included in our analysis (Figure 3). In contrast, the BRL2 fern clade was present in all ferns in our analysis, including the heterosporous species. Expression of the CtBRL2s under the control of the Arabidopsis BRI1 promoter could not complement even a weak *bri1* allele (Zheng et al. 2022). However, fusion proteins of the BRI1 extracellular receptor domain with a CtBRL2 cytoplasmic domain could complement a *bri1* allele and activate downstream events in BR signaling (Zheng et al. 2022). This raises the alternative possibility that, in ferns, the larger BRL2 clade functions in BR perception for growth and elongation in the sporophyte.

The chemical composition of A_CE_ is unknown, but it was hypothesized to be a GA-derived signaling molecule. Support for this hypothesis comes from three primary lines of evidence. First, when GA biosynthesis is blocked with a chemical inhibitor, the proportion of gametophytes that develop as males decreases in a population (Warne and Hickok 1989). Second, ABA blocks the A_CE_ response and is a well-known GA antagonist in angiosperms. Third, the B type antheridiogens of *Anemia* and *Lygodium* (Schizaeales) are GAs (Corey et al. 1986; Takeno et al. 1989; Yamauchi et al. 1996; Yamane 1998; Kurumatani et al. 2001). However, the discovery of *HER7*, a gene evolutionarily related to the BR receptor BRI1, places the above evidence in a new context. Complex interactions between BR, GA, and ABA are well documented in angiosperms (Zhang et al. 2009; Cai et al. 2014; Hu and Yu 2014; Wang et al. 2018). GA and BR gate each other’s developmental effects in maize (Best et al. 2016, 2017; Best and Dilkes 2023; Kaur et al. 2024). If similar interactions affect sexual development in Ceratopteris, a block in BR signaling in *her7* mutants could prevent A_CE_ signaling if A_CE_ was GA-derived. Likewise, inhibition of GA signaling by inhibitor treatment could impact sexual development if the A_CE_ pheromone is BR-derived. There are other alternative ways in which BRs could be involved in sex determination in Ceratopteris. For example, A_CE_ may yet be a GA (or some other signaling molecule), while BR signaling is still required for male development. The evolutionary distance between the Schizaeales and Ceratopteris, and apparent independent evolution of environmental sex determination in the two lineages, indicate that B and C type antheridiogens need not share a chemical identity (Hornych et al. 2021). Our molecular identification of the gene encoding HER7 raises the intriguing possibility that A_CE_ is a BR and that HER7 is involved in the direct perception of A_CE_. If so, the exploration of BR chemical diversity in Ceratopteris, and gametophytes in particular, should result in the discovery of both A_CE_ as well as the ligands for the CtBRL2s.

The likely presence of multiple unknown BR and GA compounds in Ceratopteris as well as the multi functionality of these signaling molecules means that it is difficult to draw conclusions from the gene expression data alone as to the function of BR and GA in sex determination. Genes involved in BR and GA biosynthesis and signaling are both active in Ceratopteris gametophytes (Figure 6). Generally, BR biosynthetic genes were more highly expressed in hermaphrodites when compared with males (Figure 6). Future work is needed to determine the chemical composition of A_CE_.

**Figure 6:**
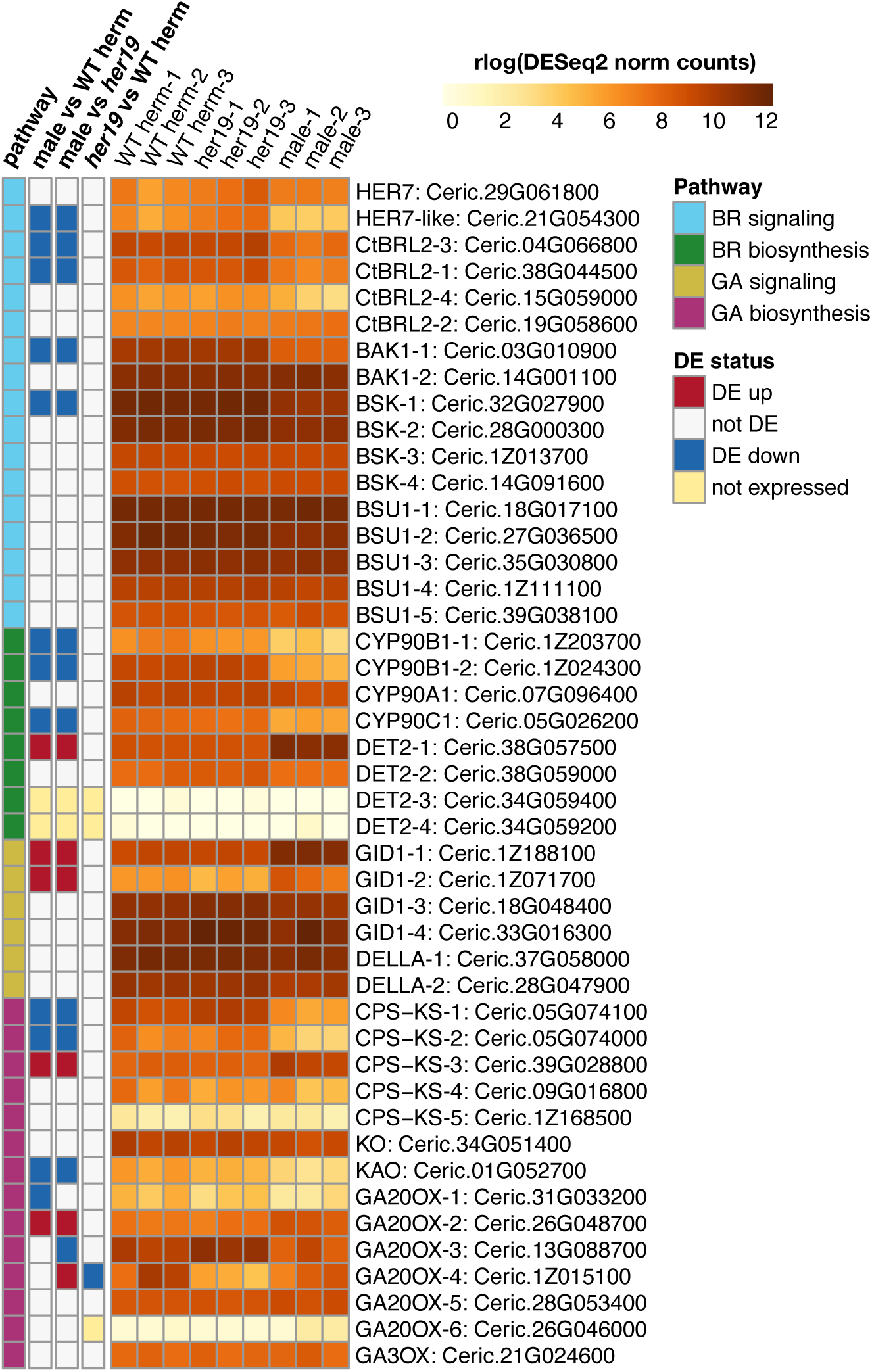
Differential expression of BR and GA signaling and biosynthesis genes.

Fortunately, a wealth of genetic work has been performed on the sex determination system of Ceratopteris and can be used to investigate several remaining open questions. Does HER7 perceive A_CE_ and if so, what is A_CE_? Are both BR and GA required for sex determination in Ceratopteris? Additionally, gene expression data indicates that *HER7* levels are similar in both male and hermaphrodite gametophytes. Does this mean that it has a different function between male and hermaphrodite gametophytes? If so, does this mean the signaling cascade of HER7 is regulated differently between the sexes? Besides the mutants used in this study, there are at least three additional *her* loci that result in an identical phenotype to the mutants present in this study (Warne et al. 1988; Eberle and Banks 1996). Additionally, there are many published sex determination mutants that we have yet to molecularly identify. These include the *fem* and *not* loci, mutants of which do not develop antheridia; the *man* locus, whose mutant develops an overabundance of antheridia; and at least two *tra* loci, that produce mutants that are always male (Banks 1994; Eberle and Banks 1996; Strain et al. 2001). Each of these loci are involved in a different part of sex determination and have stereotypically epistatic interactions with other members of the pathway (Eberle and Banks 1996). The collective knowledge of their molecular identities would provide a detailed understanding of Ceratopteris sex determination. With the advent of modern sequencing and genomic tools, this goal is now achievable.

## METHODS

### Plant growth and sample collection for BSR-Seq

Ceratopteris gametophytes were grown in either FM (fern media) or CFM (conditioned fern media; *i.e.,* FM supplemented with A_CE_ to induce male development) according to Banks et al. (1993). Agar-solidified FM/CFM was prepared with 0.7% agar (A1296, Sigma-Aldrich, St. Louis, MO, USA). FM consisted of 7 g/L of Murashige & Skoog Basal Medium with Vitamins at 6.5 pH (M519, PhytoTech Labs, Lenexa, KS, USA). For making CFM, A_CE_ was obtained as a crude aqueous filtrate of liquid FM that had supported the growth of gametophytes from *her* spores (50 mg spores/liter FM) for 28 days on a shaking incubator. Each batch of CFM was assayed for A_CE_ activity (>95% males on test plates of Hn-n wild type gametophytes) according to Banks et al. (1993). All spores were surface sterilized in a 1:3 bleach dilution with a drop of tween prior to plating.

All antheridiogen-insensitive *hermaphroditic* (*her*) mutants used in our experiments were isolated by Banks (1994) using EMS mutagenesis screens in the Hn-n genetic background. The wild type accession ΦN8 from Nicaragua (Hickok et al. 1995) was crossed with *her7-14* and *her7-19* for bulk segregant analysis. Crosses were performed according to Eberle and Banks (1996). Briefly, 9-12-day-old individual *her* hermaphrodites were placed in microtiter wells of 24-well plates with agar solidified FM. For acquiring male gamete donors, ΦN8 spores were plated on CFM and maintained at 29 °C. At 11-13 days after spore inoculation, room temperature water was added to the ΦN8 males, causing the sperm to be released. Drops of water containing ΦN8 sperm were added to each individual *her* hermaphrodite. Sperm swarming the archegonia was confirmed visually. Conditions used for growing Ceratopteris sporophytes are described in Banks (1994). Hybrid *her7-14* x ΦN8 and *her7-19* x ΦN8 spore progeny were plated on CFM and grown at 29 °C. After 10 days, four pooled samples were collected by manual picking of n ≈ 1000 individual gametophytes: M14 (*her7-14* x ΦN8 males), H14 (*her7-14* x ΦN8 hermaphrodites), M19 (*her7-19* x ΦN8 males), and H19 (*her7-19* x ΦN8 hermaphrodites). Samples were frozen in liquid nitrogen and stored at -80°C.

Wild type Hn-n and ΦN8 spores were grown in the same conditions as described above and plated on FM. After 10 days, pools of male and hermaphrodite gametophytes were collected by manual picking of n ≈ 1000 individual gametophytes for each accession.

Previously published Ceratopteris RNA-seq data (Geng et al. 2018) was downloaded from NCBI. RNA-seq samples and data availability are listed in Table S5.

### RNA extraction, sequencing, and processing

Total RNA was extracted with a Quick-RNA Plant miniprep kit (Zymo Research, Irvine, CA, USA) using a Disruptor Genie (Scientific Industries, Bohemia, NY, USA) for tissue disruption. RNA was quantified using a Qubit 4 fluorometer (ThermoFisher Scientific, Waltham, MA, USA). RNA integrity (RIN) was assessed using a TapeStation 4150 (Agilent, Santa Clara, CA, USA), and samples with RIN ≥ 5.9 were analyzed for expression. For Hn-n and ΦN8, total RNA from hermaphrodite and male samples were combined to create a single reference sample per wild type accession (Table S5).

Samples were shipped to Novogene Corporation Inc. (Sacramento, CA, USA) for library construction (NEBNext Ultra RNA Library Prep Kit, New England Biolabs, Ipswich, MA, USA) from 1 µg total RNA and Illumina sequencing using the NovaSeq 6000 Sequencing System. Paired-end (2x150 bp) reads were processed with fastp v0.20.1 (Chen et al. 2018) to trim reads with no signal, remove adaptor sequence, and correct for mismatches in overlapping reads. Read quality was checked with MultiQC v1.9 (Ewels et al. 2016).

### Variant calling

To identify high-quality single nucleotide polymorphisms (SNPs) distinguishing the ΦN8 background and Hn-n reference, ΦN8 and Hn-n reads were aligned to the Hn-n reference genome (Marchant et al. 2022) using STAR v2.7.10 (Dobin et al. 2013) with default parameters. The percentage of uniquely mapped reads was between 88-95% for all samples (Table S5). Variant calling was performed using two pipelines. First, mapped reads were processed with GATK v4.2.6.1 (Brouard et al. 2019) and Picard v2.27.1 using MarkDuplicates and SplitNCigars from GATK’s large genome RNA-seq processing pipeline to process duplicate reads and spliced reads. Next, processed reads were annotated using AddReadGroups, and polymorphisms were identified using HaplotypeCaller. For the second pipeline, variants were identified using samtools v1.11 and bcftools v1.11 (Danecek et al. 2021). STAR aligned reads were preprocessed using sort, fixmate, and markdup followed by variant calling with mpileup. Only SNPs recovered by both pipelines were retained for further analysis using bcftools.

Additionally, quality filtration was performed using SnpSift v4.3.1t (Cingolani et al. 2012), which removed indels, excluded non-biallelic sites, required a quality score > 20, read depth > 4, and stranded depth > 2. Variants on unplaced scaffolds were excluded. Lastly, ΦN8 and Hn-n SNPs were intersected and only SNPs unique to ΦN8 were retained.

#### Bulked segregant analysis

To map the location of the *her7-14* and *her7-19* mutations, SNPs were identified in the *her* x ΦN8 RNAseq samples (M14, H14, M19, and H19) following the same variant calling pipelines as described above, and variant calls in hybrid samples were intersected with the ΦN8 variant list. Read depths for the alternative (i.e., ΦN8-derived) allele were used to calculate ΦN8 allele frequencies in 16-Mbp windows across the entire genome, and an allele frequency ratio summarizing the distribution of the ΦN8 genotype was calculated using the following formula:

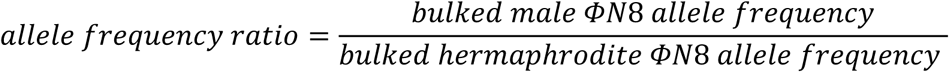

Variants in the *her7-19* RNA-seq dataset (Geng et al. 2021) were identified following the same variant calling pipelines as described above, and Ensembl’s Variant Effect Predictor v108.2 (McLaren et al. 2016) was used to flag variants with moderate to high effect on the predicted protein products.

#### PCR amplification

The *HER7* sequence in wild type (Hn-n and ΦN8) and *her* mutants was determined by sequencing PCR amplicons. Genomic DNA was extracted from 12-day old gametophytes using Quick-DNA Plant/Seed Miniprep Kit (Zymo Research, Irvine, CA, USA). Primers were designed using SnapGene with elongation factor 1 (Ceric.09G081400) used as a positive control (Table S6). PCR amplification was performed in 50 μL reactions as follows: 25 μL 2x Q5 High Fidelity master mix (New England Biolabs Inc, Ipswich, MA, USA), 5 μL each of the forward and reverse primers (500 nM final concentration), and approximately 100 ng of template DNA. Cycling parameters included an initial denaturation step at 98°C for 30 sec followed by 25-35 cycles of 98°C for 10 sec, 71°C for 20 sec, 72°C for 2 min 20 sec and a final elongation step at 72°C for 2 min. PCR products were visualized on an agarose gel to confirm expected DNA size range and purified using the ChargeSwitch-Pro PCR Cleanup Kit (Invitrogen, Waltham, MA, USA). PCR amplicons were sequenced using the WideSeq service provided by the Purdue Genomics Core Facility, which generated paired-end reads (2 x 250 bp) using an Illumina MiSeq sequencing machine. Reads from each sample were processed to remove adapters and poor-quality bases using Trimmomatic v0.39 (Bolger et al. 2014) with the following settings: PE LEADING:20, TRAILING:20, MINLEN:30, and ILLUMINACLIP of adapters set to 2:40:10. Filtered paired reads were aligned to the genome with BWA v 0.7.17-r1188 (Li and Durbin 2009), and variants were identified with HaplotypeCaller as described above. The detected variants were manually inspected by visualizing each sample in IGV version 2.8.2 (Robinson et al. 2011) using the sorted BAM files.

#### HER7 phylogeny

Homology between the predicted proteomes of Ceratopteris and 37 other plant species (Table S7) was determined with OrthoFinder v2.4.1 (Emms and Kelly 2019) with sequence similarity searches performed by DIAMOND v2.0.8.146 (Buchfink et al. 2015). The orthogroup containing *HER7* (Ceric.29G061800) was identified, and the corresponding amino acid sequences were aligned with MAFFT v7.471 using the E-INS-i iterative refinement method (Katoh and Standley 2013). An initial maximum likelihood (ML) phylogeny was constructed with IQ-TREE v1.6.12 (Nguyen et al. 2015) using the built in ModelFinder search (Kalyaanamoorthy et al. 2017) to determine the best-fit amino acid substitution model (JTT+R6). GeneRax v2.0.4 (Morel et al. 2020) was used to optimize the branching order of the resulting ML gene family tree based on the species tree (Figure 3B) and a JTT+G substitution model. The root of the HER7 phylogeny as well as patterns of gene duplication and loss were also inferred from the GeneRax output. A topology test was performed in IQ-TREE that compared the log-likelihoods of three trees: 1) the initial ML tree, 2) the optimized GeneRax tree, and 3) a constrained tree where all fern sequences were forced to be monophyletic. The same ML analysis was repeated for orthogroups containing genes for BR biosynthesis; however, GeneRax reconciliation analyses were not performed, and ML trees were manually rooted based on the species tree. All trees were visualized using the ETE v3 phylogenetic toolkit (Huerta-Cepas et al. 2016).

#### Plasmid construction, transient expression, and confocal imaging

The *HER7* coding sequence (CDS) was amplified from the genomic DNA of wild-type Ceratopteris (Hn-n) using the primers provided in Table S6 and cloned into the pENTR vector through enzyme digestion and ligation. Subsequently, the *HER7* coding sequence was introduced into the pMDC43 vector (Curtis and Grossniklaus 2003) to generate a *35S::GFP-HER7* expression cassette. Transient expression in *Nicotiana benthamiana* leaves was performed through *Agrobacterium*-mediated infiltration, as described previously (Han et al. 2020). Specifically, *N. benthamiana* plants were grown under short-day conditions (8h-light/16h-dark) at room temperature, and plants with four leaves were used for the assay. Each construct was transformed into *Agrobacterium tumefaciens* strain GV3101 through electroporation. Cultures of agrobacteria containing *35S::GFP-HER7* (in pMDC43) or the P19 silencing suppressor were resuspended and mixed in a solution of 10 mM MgCl_2_ and 150 μM acetosyringone and then infiltrated into *N. benthamiana* leaves. The culture mix of *35S::GFP* (in a pMDC43 vector with the gateway cassette removed) and P19 was prepared using the same procedure as a control. At least three individual plants were infiltrated with either *35S::GFP-HER7* or *35S::GFP*.

Two days after infiltration, the samples were imaged using a Zeiss LSM880 confocal laser microscope with a Plan-Apochromat 10x/0.45 objective lens and a 1.0 μm scanning interval. GFP was excited with a 488 nm laser line, and emission was collected within the 490-544 nm range. DIC imaging was performed using the 488 nm laser line, with signals collected through the T-PMT (transmitted photomultiplier tubes) module to visualize cell outlines. Maximum-intensity Z projection views of the confocal stacks were generated using Fiji/Image J software.

#### Gene expression quantification and differential expression analysis

Wild type Hn-n spores were grown on FM media. After 12 days, hermaphrodite and male gametophytes were picked and collected at a final weight of ∼0.1g per sample. At the same time, *her7-19* spores were plated and grown on CFM and sampled at 12 days. All samples were collected in triplicate. RNA extraction, quality control, and sequencing was performed as described for the BSR-Seq analysis above. Quantification of gene expression was performed using Kallisto v0.48.0 (Bray et al. 2016). The Kallisto index was built using all Ceratopteris predicted transcripts with the default k-mer size of 31.

For the differential expression analysis, raw gene counts from Kallisto were passed to DESeq2 v.1.44.0 (Love et al. 2014) using tximport v1.32.0 (Soneson et al. 2016), and tests for significance were conducted using a Wald test using the DESeq function. The Benjamini-Hochberg (BH) multiple test correction (Benjamini and Hochberg 1995) was used to adjust p-values, and genes were considered significantly differentially expressed if adjusted p-value < 0.1 and |LFC| > 1.0. Three independent comparisons were performed: male vs hermaphrodite; male vs *her7-19*; hermaphrodite vs *her7-19*.

KEGG annotations were assigned using eggNOG-mapper (Cantalapiedra et al. 2021). Tests for enrichment of higher-level functional categories were performed using the KEGG PATHWAY metabolic hierarchy downloaded via the KEGG API (Kanehisa et al. 2014). Hypergeometric tests were performed in python using the SciPy library hypergeom (Virtanen et al. 2020) and p-values were adjusted for multiple comparisons using the StatsModels library multitest (Seabold and Perktold 2010) with the BH method (Benjamini and Hochberg 1995).

## ACKNOWLEDGMENTS

We thank Dr. Jo Ann Banks for mentoring and assisting with Ceratopteris genetics and propagation. This work was conducted in part using the resources of the Rosen Center for Advanced Computing at Purdue University. This work was supported by Purdue University through a research seed grant provided by the Center for Plant Biology in the College of Agriculture to BPD and JHW. This work was also supported in part by the National Science Foundation under grants DEB-1831493 to JHW and IOS-1931114 to YZ.

## DATA AVAILABILITY

Orthogroups, multiple sequence alignments, tree files, and other related data files are available through FigShare (10.6084/m9.figshare.27169293). Scripts are available through GitHub (https://github.com/WisecaverLab/ceratopteris_her7). Raw sequencing reads have been deposited in the Sequence Read Archive database under BioProject access number PRJNA1167630.

## References

Achard P, Cheng H, De Grauwe L, Decat J, Schoutteten H, Moritz T, Van Der Straeten D, Peng J, and Harberd NP. Integration of Plant Responses to Environmentally Activated Phytohormonal Signals. Science. 2006:311(5757):91–94. 10.1126/science.1118642

Banks JA. Sex-determining genes in the homosporous fern Ceratopteris. Development. 1994:120(7):1949–1958.

Banks JA. The TRANSFORMER Genes of the Fern Ceratopteris Simultaneously Promote Meristem and Archegonia Development and Repress Antheridia Development in the Developing Gametophyte. Genetics. 1997:147(4):1885– 1897. 10.1093/genetics/147.4.1885

Banks JA, Hickok L, and Webb MA. The Programming of Sexual Phenotype in the Homosporous Fern Ceratopteris richardii. International Journal of Plant Sciences. 1993:154(4):522–534. 10.1086/297135

Benjamini Y and Hochberg Y. Controlling the false discovery rate: a practical and powerful approach to multiple testing. Journal of the Royal Statistical Society Series B (Methodological). 1995:57(1):289–300. 10.1111/j.2517-6161.1995.tb02031.x

Best N and Dilkes B. Genetic evidence that brassinosteroids suppress pistils in the maize tassel independent of the jasmonic acid pathway. Plant Direct. 2023:7(7):e501. 10.1002/pld3.501

Best NB, Hartwig T, Budka J, Fujioka S, Johal G, Schulz B, and Dilkes BP. nana plant2 Encodes a Maize Ortholog of the Arabidopsis Brassinosteroid Biosynthesis Gene DWARF1, Identifying Developmental Interactions between Brassinosteroids and Gibberellins. Plant Physiology. 2016:171(4):2633–2647. 10.1104/pp.16.00399

Best NB, Johal G, and Dilkes BP. Phytohormone inhibitor treatments phenocopy brassinosteroid–gibberellin dwarf mutant interactions in maize. Plant Direct. 2017:1(2). 10.1002/pld3.9

Bolger AM, Lohse M, and Usadel B. Trimmomatic: a flexible trimmer for Illumina sequence data. Bioinformatics (Oxford, England). 2014:30(15):2114–2120. 10.1093/bioinformatics/btu170

Bray NL, Pimentel H, Melsted P, and Pachter L. Near-optimal probabilistic RNA-seq quantification. Nature Biotechnology. 2016:34(5):525–527. 10.1038/nbt.3519

Brouard J-S, Schenkel F, Marete A, and Bissonnette N. The GATK joint genotyping workflow is appropriate for calling variants in RNA-seq experiments. Journal of Animal Science and Biotechnology. 2019:10(1):44. 10.1186/s40104-019-0359-0

Buchfink B, Xie C, and Huson DH. Fast and sensitive protein alignment using DIAMOND. Nature methods. 2015:12(1):59–60. 10.1038/nmeth.3176

Cai Z, Liu J, Wang H, Yang C, Chen Y, Li Y, Pan S, Dong R, Tang G, Barajas-Lopez J de D, et al. GSK3-like kinases positively modulate abscisic acid signaling through phosphorylating subgroup III SnRK2s in Arabidopsis. Proceedings of the National Academy of Sciences. 2014:111(26):9651–9656. 10.1073/pnas.1316717111

Cantalapiedra CP, Hernández-Plaza A, Letunic I, Bork P, and Huerta-Cepas J. eggNOG-mapper v2: Functional annotation, orthology assignments, and domain prediction at the metagenomic scale. Molecular Biology and Evolution. 2021:38(12):5825–5829. 10.1093/molbev/msab293

Chatterjee A and Roux SJ. Ceratopteris richardii: A Productive Model for Revealing Secrets of Signaling and Development. J Plant Growth Regul. 2000:19(3):284–289. 10.1007/s003440000032

Chen S, Zhou Y, Chen Y, and Gu J. fastp: an ultra-fast all-in-one FASTQ preprocessor. Bioinformatics. 2018:34(17):i884–i890. 10.1093/bioinformatics/bty560

Cheon J, Fujioka S, Dilkes BP, and Choe S. Brassinosteroids Regulate Plant Growth through Distinct Signaling Pathways in Selaginella and Arabidopsis. PLOS ONE. 2013:8(12):e81938. 10.1371/journal.pone.0081938

Cheon J, Park S-Y, Schulz B, and Choe S. Arabidopsis brassinosteroid biosynthetic mutant dwarf7-1exhibits slower rates of cell division and shoot induction. BMC Plant Biol. 2010:10(1):270. 10.1186/1471-2229-10-270

Chung Y and Choe S. The Regulation of Brassinosteroid Biosynthesis in Arabidopsis. Critical Reviews in Plant Sciences. 2013:32(6):396–410. 10.1080/07352689.2013.797856

Cingolani P, Platts A, Wang LL, Coon M, Nguyen T, Wang L, Land SJ, Lu X, and Ruden DM. A program for annotating and predicting the effects of single nucleotide polymorphisms, SnpEff: SNPs in the genome of Drosophila melanogaster strain w ^1118^ ; iso-2; iso-3. Fly. 2012:6(2):80–92. 10.4161/fly.19695

Corey EJ, Myers AG, Takahashi N, Yamane H, and Schraudolf H. Constitution of antheridium-inducing factor of *anemia phyllitidis*. Tetrahedron Letters. 1986:27(42):5083–5084. 10.1016/S0040-4039(00)85138-2

Curtis MD and Grossniklaus U. A Gateway Cloning Vector Set for High-Throughput Functional Analysis of Genes in Planta. Plant Physiology. 2003:133(2):462–469. 10.1104/pp.103.027979

Danecek P, Bonfield JK, Liddle J, Marshall J, Ohan V, Pollard MO, Whitwham A, Keane T, McCarthy SA, Davies RM, et al. Twelve years of SAMtools and BCFtools. Gigascience. 2021:10(2). 10.1093/gigascience/giab008

Dobin A, Davis CA, Schlesinger F, Drenkow J, Zaleski C, Jha S, Batut P, Chaisson M, and Gingeras TR. STAR: ultrafast universal RNA-seq aligner. Bioinformatics. 2013:29(1):15–21. 10.1093/bioinformatics/bts635

Döpp W. Eine die Antheridienbildung bei Farnen fördernde Substanz in den Prothallien von Pteridium aquilinum (L.) Kuhn. Berichte der Deutschen Botanischen Gesellschaft. 1950:63(5):139–147. 10.1111/j.1438-8677.1951.tb01498.x

Eberle J, Nemacheck J, Wen C-K, Hasebe M, and Banks JA. Ceratopteris: A Model System for Studying Sex-Determining Mechanisms in Plants. International Journal of Plant Sciences. 1995:156(3):359–366.

Eberle JR and Banks JA. Genetic Interactions Among Sex-Determining Genes in the Fern Ceratopteris richardii. Genetics. 1996:142(3):973–985. 10.1093/genetics/142.3.973

Emms DM and Kelly S. OrthoFinder: Phylogenetic orthology inference for comparative genomics. Genome Biology. 2019:(20):1. 10.1186/s13059-019-1832-y

Ewels P, Magnusson M, Lundin S, and Käller M. MultiQC: summarize analysis results for multiple tools and samples in a single report. Bioinformatics. 2016:32(19):3047–3048. 10.1093/bioinformatics/btw354

Fernández H, Grossmann J, Gagliardini V, Feito I, Rivera A, Rodríguez L, Quintanilla LG, Quesada V, Cañal MJ, and Grossniklaus U. Sexual and Apogamous Species of Woodferns Show Different Protein and Phytohormone Profiles. Front Plant Sci. 2021:12:718932. 10.3389/fpls.2021.718932

Finch-Savage WE and Leubner-Metzger G. Seed dormancy and the control of germination. New Phytol. 2006:171(3):501–523. 10.1111/j.1469-8137.2006.01787.x

Fujioka S, Li J, Choi YH, Seto H, Takatsuto S, Noguchi T, Watanabe T, Kuriyama H, Yokota T, Chory J, et al. The Arabidopsis deetiolated2 mutant is blocked early in brassinosteroid biosynthesis. Plant Cell. 1997:9(11):1951–1962. 10.1105/tpc.9.11.1951

Furuya T, Ohashi-Ito K, and Kondo Y. Multiple Roles of Brassinosteroid Signaling in Vascular Development. Plant and Cell Physiology. 2024:pcae037. 10.1093/pcp/pcae037

Geng S, Miyagi A, and Umen JG. Evolutionary divergence of the sex-determining gene MID uncoupled from the transition to anisogamy in volvocine algae. Development. 2018:145(7):dev162537. 10.1242/dev.162537

Geng Y, Cai C, McAdam SAM, Banks JA, Wisecaver JH, and Zhou Y. A De Novo Transcriptome Assembly of Ceratopteris richardii Provides Insights into the Evolutionary Dynamics of Complex Gene Families in Land Plants. Genome Biology and Evolution. 2021:13(3):evab042. 10.1093/gbe/evab042

Han H, Yan A, Li L, Zhu Y, Feng B, Liu X, and Zhou Y. A signal cascade originated from epidermis defines apical-basal patterning of Arabidopsis shoot apical meristems. Nat Commun. 2020:11(1):1214. 10.1038/s41467-020-14989-4

Hartwig T, Chuck GS, Fujioka S, Klempien A, Weizbauer R, Potluri DPV, Choe S, Johal GS, and Schulz B. Brassinosteroid control of sex determination in maize. Proc Natl Acad Sci U S A. 2011:108(49):19814–19819. 10.1073/pnas.1108359108

He Z, Wang Z-Y, Li J, Zhu Q, Lamb C, Ronald P, and Chory J. Perception of Brassinosteroids by the Extracellular Domain of the Receptor Kinase BRI1. Science. 2000:288(5475):2360–2363. 10.1126/science.288.5475.2360

Hedden P and Phillips AL. Gibberellin metabolism: new insights revealed by the genes. Trends in Plant Science. 2000:5(12):523–530. 10.1016/S1360-1385(00)01790-8

Helliwell CA, Chandler PM, Poole A, Dennis ES, and Peacock WJ. The CYP88A cytochrome P450, ent-kaurenoic acid oxidase, catalyzes three steps of the gibberellin biosynthesis pathway. Proceedings of the National Academy of Sciences. 2001:98(4):2065–2070. 10.1073/pnas.98.4.2065

Helliwell CA, Sheldon CC, Olive MR, Walker AR, Zeevaart JAD, Peacock WJ, and Dennis ES. Cloning of the Arabidopsis ent-kaurene oxidase gene GA3. Proceedings of the National Academy of Sciences. 1998:95(15):9019–9024. 10.1073/pnas.95.15.9019

Hickok LG. Abscisic acid blocks antheridiogen-induced antheridium formation in gametophytes of the fern Ceratopteris. Can J Bot. 1983:61(3):888–892. 10.1139/b83-098

Hickok LG, Warne TR, and Frisbourg RS. The Biology of the Fern Ceratopteris and Its Use as a Model System. International Journal of Plant Sciences. 1995:156:332– 345.

Hickok LG, Warne TR, and Slocum MK. Ceratopteris Richardii: Applications for Experimental Plant Biology. American Journal of Botany. 1987:74(8):1304–1316. 10.1002/j.1537-2197.1987.tb08743.x

Hornych O, Testo WL, Sessa EB, Watkins Jr. JE, Campany CE, Pittermann J, and Ekrt L. Insights into the evolutionary history and widespread occurrence of antheridiogen systems in ferns. New Phytologist. 2021:229(1):607–619. 10.1111/nph.16836

Hu Y and Yu D. BRASSINOSTEROID INSENSITIVE2 Interacts with ABSCISIC ACID INSENSITIVE5 to Mediate the Antagonism of Brassinosteroids to Abscisic Acid during Seed Germination in Arabidopsis. The Plant Cell. 2014:26(11):4394–4408. 10.1105/tpc.114.130849

Huerta-Cepas J, Serra F, and Bork P. ETE 3: Reconstruction, Analysis, and Visualization of Phylogenomic Data. Molecular Biology and Evolution. 2016:33(6):1635–1638. 10.1093/molbev/msw046

Hussain A and Peng J. DELLA Proteins and GA Signalling in Arabidopsis. J Plant Growth Regul. 2003:22(2):134–140. 10.1007/s00344-003-0028-5

Kalyaanamoorthy S, Minh BQ, Wong TKF, Von Haeseler A, and Jermiin LS. ModelFinder: Fast model selection for accurate phylogenetic estimates. Nature Methods. 2017. 10.1038/nmeth.4285

Kanehisa M, Goto S, Sato Y, Kawashima M, Furumichi M, and Tanabe M. Data, information, knowledge and principle: back to metabolism in KEGG. Nucleic Acids Research. 2014:42(Database issue):D199-205.

Katoh K and Standley DM. MAFFT multiple sequence alignment software version 7: Improvements in performance and usability. Molecular Biology and Evolution. 2013:30(4):772–780. 10.1093/molbev/mst010

Kaur A, Best NB, Hartwig T, Budka J, Khangura RS, McKenzie S, Aragón-Raygoza A, Strable J, Schulz B, and Dilkes BP. A maize semi-dwarf mutant reveals a GRAS transcription factor involved in brassinosteroid signaling. Plant Physiology. 2024:195(4):3072–3096. 10.1093/plphys/kiae147

Kim BK, Fujioka S, Takatsuto S, Tsujimoto M, and Choe S. Castasterone is a likely end product of brassinosteroid biosynthetic pathway in rice. Biochem Biophys Res Commun. 2008:374(4):614–619. 10.1016/j.bbrc.2008.07.073

Kim T-W, Michniewicz M, Bergmann DC, and Wang Z-Y. Brassinosteroid regulates stomatal development by GSK3-mediated inhibition of a MAPK pathway. Nature. 2012:482(7385):419–422. 10.1038/nature10794

Kurumatani M, Yagi K, Murata T, Tezuka M, Mander LN, Nishiyama M, and Yama H. Isolation and identification of antheridiogens in the ferns, Lygodium microphyllum and Lygodium reticulatum. Biosci Biotechnol Biochem. 2001:65(10):2311–2314. 10.1271/bbb.65.2311

Li H and Durbin R. Fast and accurate short read alignment with Burrows–Wheeler transform. Bioinformatics. 2009:25(14):1754–1760. 10.1093/bioinformatics/btp324

Li J, Wen J, Lease KA, Doke JT, Tax FE, and Walker JC. BAK1, an Arabidopsis LRR receptor-like protein kinase, interacts with BRI1 and modulates brassinosteroid signaling. Cell. 2002:110(2):213–222. 10.1016/s0092-8674(02)00812-7

Love MI, Huber W, and Anders S. Moderated estimation of fold change and dispersion for RNA-seq data with DESeq2. Genome Biology. 2014:15(12):1–21. 10.1186/s13059-014-0550-8

Makarevitch I, Thompson A, Muehlbauer GJ, and Springer NM. Brd1 gene in maize encodes a brassinosteroid C-6 oxidase. PLoS One. 2012:7(1):e30798. 10.1371/journal.pone.0030798

Marchant DB, Chen G, Cai S, Chen F, Schafran P, Jenkins J, Shu S, Plott C, Webber J, Lovell JT, et al. Dynamic genome evolution in a model fern. Nat Plants. 2022:8(9):1038–1051. 10.1038/s41477-022-01226-7

McLaren W, Gil L, Hunt SE, Riat HS, Ritchie GRS, Thormann A, Flicek P, and Cunningham F. The Ensembl Variant Effect Predictor. Genome Biology. 2016:17(1):122. 10.1186/s13059-016-0974-4

Moran RC. A natural history of ferns (Timber Press: Portland).

Morel B, Kozlov AM, Stamatakis A, and Szöllősi GJ. GeneRax: A Tool for Species-Tree- Aware Maximum Likelihood-Based Gene Family Tree Inference under Gene Duplication, Transfer, and Loss. Molecular Biology and Evolution. 2020:37(9):2763–2774. 10.1093/molbev/msaa141

Nguyen LT, Schmidt HA, Von Haeseler A, and Minh BQ. IQ-TREE: A fast and effective stochastic algorithm for estimating maximum-likelihood phylogenies. Molecular Biology and Evolution. 2015:32(1):268–274. 10.1093/molbev/msu300

Noguchi T, Fujioka S, Choe S, Takatsuto S, Yoshida S, Yuan H, Feldmann KA, and Tax FE. Brassinosteroid-Insensitive Dwarf Mutants of Arabidopsis Accumulate Brassinosteroids1. Plant Physiology. 1999:121(3):743–752. 10.1104/pp.121.3.743

PPG. A community-derived classification for extant lycophytes and ferns. Journal of Systematics and Evolution. 2016:54(6):563–603. 10.1111/jse.12229

Robinson JT, Thorvaldsdóttir H, Winckler W, Guttman M, Lander ES, Getz G, and Mesirov JP. Integrative genomics viewer. Nat Biotechnol. 2011:29(1):24–26. 10.1038/nbt.1754

Ryu H, Kim K, Cho H, and Hwang I. Predominant actions of cytosolic BSU1 and nuclear BIN2 regulate subcellular localization of BES1 in brassinosteroid signaling. Mol Cells. 2010:29(3):291–296. 10.1007/s10059-010-0034-y

Schneller JJ. Antheridiogens. . In. Biology and Evolution of Ferns and Lycophytes, CH Haufler and TA Ranker, eds. (Cambridge University Press: Cambridge), pp. 134–158. 10.1017/CBO9780511541827.006

Seabold S and Perktold J. Statsmodels: Econometric and Statistical Modeling with Python. Proceedings of the 9th Python in Science Conference. 2010:92–96. 10.25080/Majora-92bf1922-011

She J, Han Z, Kim T-W, Wang J, Cheng W, Chang J, Shi S, Wang J, Yang M, Wang Z-Y, et al. Structural insight into brassinosteroid perception by BRI1. Nature. 2011:474(7352):472–476. 10.1038/nature10178

Shi H, Yan H, Li J, and Tang D. BSK1, a receptor-like cytoplasmic kinase, involved in both BR signaling and innate immunity in Arabidopsis. Plant Signal Behav. 2013:8(8):e24996. 10.4161/psb.24996

Soneson C, Love MI, and Robinson MD. Differential analyses for RNA-seq: transcript- level estimates improve gene-level inferences. 2016. 10.12688/f1000research.7563.2

Strain E, Hass B, and Banks JA. Characterization of Mutations That Feminize Gametophytes of the Fern Ceratopteris. Genetics. 2001:159(3):1271–1281. 10.1093/genetics/159.3.1271

Sun C, Yan K, Han J-T, Tao L, Lv M-H, Shi T, He Y-X, Wierzba M, Tax FE, and Li J. Scanning for New BRI1 Mutations via TILLING Analysis. Plant Physiology. 2017:174(3):1881–1896. 10.1104/pp.17.00118

Sun TP and Kamiya Y. The Arabidopsis GA1 locus encodes the cyclase ent-kaurene synthetase A of gibberellin biosynthesis. The Plant Cell. 1994:6(10):1509–1518. 10.1105/tpc.6.10.1509

Takeno K, Yamane H, Yamauchi T, Takahashi N, Furber M, and Mander LN. Biological Activities of the Methyl Ester of Gibberellin A73, a Novel and Principal Antheridiogen in Lygodium japonicum. Plant and Cell Physiology. 1989:30(2):201–205. 10.1093/oxfordjournals.pcp.a077730

Thomas SG, Phillips AL, and Hedden P. Molecular cloning and functional expression of gibberellin 2- oxidases, multifunctional enzymes involved in gibberellin deactivation. Proceedings of the National Academy of Sciences. 1999:96(8):4698–4703. 10.1073/pnas.96.8.4698

Ueguchi-Tanaka M, Ashikari M, Nakajima M, Itoh H, Katoh E, Kobayashi M, Chow T, Hsing YC, Kitano H, Yamaguchi I, et al. GIBBERELLIN INSENSITIVE DWARF1 encodes a soluble receptor for gibberellin. Nature. 2005:437(7059):693–698. 10.1038/nature04028

Virtanen P, Gommers R, Oliphant TE, Haberland M, Reddy T, Cournapeau D, Burovski E, Peterson P, Weckesser W, Bright J, et al. SciPy 1.0: fundamental algorithms for scientific computing in Python. Nat Methods. 2020:17(3):261–272. 10.1038/s41592-019-0686-2

Vriet C, Russinova E, and Reuzeau C. From Squalene to Brassinolide: The Steroid Metabolic and Signaling Pathways across the Plant Kingdom. Molecular Plant. 2013:6(6):1738–1757. 10.1093/mp/sst096

Wang H, Tang J, Liu J, Hu J, Liu J, Chen Y, Cai Z, and Wang X. Abscisic Acid Signaling Inhibits Brassinosteroid Signaling through Dampening the Dephosphorylation of BIN2 by ABI1 and ABI2. Molecular Plant. 2018:11(2):315–325. 10.1016/j.molp.2017.12.013

Wang Z-Y, Bai M-Y, Oh E, and Zhu J-Y. Brassinosteroid Signaling Network and Regulation of Photomorphogenesis. Annual Review of Genetics. 2012:46(Volume 46, 2012):701–724. 10.1146/annurev-genet-102209-163450

Warne TR and Hickok LG. Evidence for a gibberellin biosynthetic origin of ceratopteris antheridiogen. Plant Physiol. 1989:89(2):535–538. 10.1104/pp.89.2.535

Warne TR, Hickok LG, and Scott RJ. Characterization and genetic analysis of antheridiogen-insensitive mutants in the fern Ceratopteris. Botanical Journal of the Linnean Society. 1988:96(4):371–379. 10.1111/j.1095-8339.1988.tb00692.x

Yamaguchi S, Sun T, Kawaide H, and Kamiya Y. The GA2 Locus of Arabidopsis thalianaEncodes ent-Kaurene Synthase of Gibberellin Biosynthesis. Plant Physiology. 1998:116(4):1271–1278. 10.1104/pp.116.4.1271

Yamane H. Fern Antheridiogens. . In. International Review of Cytology, KW Jeon, ed. (Academic Press), pp. 1–32. 10.1016/S0074-7696(08)62177-4

Yamauchi T, Oyama N, Yamane H, Murofushi N, Schraudolf H, Pour M, Furber M, and Mander LN. Identification of Antheridiogens in Lygodium circinnatum and Lygodium flexuosum. Plant Physiol. 1996:111(3):741–745. 10.1104/pp.111.3.741

Yang J, Thames S, Best NB, Jiang H, Huang P, Dilkes BP, and Eveland AL. Brassinosteroids Modulate Meristem Fate and Differentiation of Unique Inflorescence Morphology in Setaria viridis. The Plant Cell. 2018:30(1):48–66. 10.1105/tpc.17.00816

Yokota T, Ohnishi T, Shibata K, Asahina M, Nomura T, Fujita T, Ishizaki K, and Kohchi T. Occurrence of brassinosteroids in non-flowering land plants, liverwort, moss, lycophyte and fern. Phytochemistry. 2017:136:46–55. 10.1016/j.phytochem.2016.12.020

Zhang S, Cai Z, and Wang X. The primary signaling outputs of brassinosteroids are regulated by abscisic acid signaling. Proceedings of the National Academy of Sciences. 2009:106(11):4543–4548. 10.1073/pnas.0900349106

Zheng B, Xing K, Zhang J, Liu H, Ali K, Li W, Bai Q, and Ren H. Evolutionary Analysis and Functional Identification of Ancient Brassinosteroid Receptors in Ceratopteris richardii. International Journal of Molecular Sciences. 2022:23(12):6795. 10.3390/ijms23126795

